# refineDLC: an advanced post-processing pipeline for DeepLabCut outputs

**DOI:** 10.1101/2025.04.09.648046

**Authors:** Weronika Klecel, Hadley Rahael, Samantha Ann Brooks

## Abstract

DeepLabCut has notably transformed behavioral and locomotor research by enabling precise, markerless pose estimation through advanced deep learning methods. Despite broad adoption of this approach across species and behaviors, current usage frequently overlooks the challenges of quantitative kinematic analyses due to the complexity and noise within raw model outputs. Researchers are often challenged by the computational expertise required to clean, refine, and interpret the generated data effectively. Here, we introduce refineDLC, a comprehensive post-processing pipeline explicitly designed to streamline the conversion of noisy DeepLabCut outputs into robust, analytically reliable kinematic data.

Our user-friendly pipeline incorporates critical data cleaning steps—including inversion of y-coordinates for intuitive spatial interpretation, removal of zero-value frames, and exclusion of irrelevant body part labels. Additionally, it integrates robust, dual-stage filtering methods based on likelihood scores and positional changes, significantly enhancing the accuracy and consistency of data. Finally, our approach offers multiple interpolation strategies, effectively managing missing values to maintain data continuity and integrity. We evaluated the performance and versatility of our pipeline across two distinct datasets: controlled locomotion in cattle and field-recorded, highly variable trotting horse videos.

Our results demonstrated substantial improvements in the quality and interpretability of processed outputs, transforming initially noisy and inconsistent data into physiologically meaningful kinematic patterns. RefineDLC successfully reduced data variability and eliminated false-positive labeling errors, offering reliable and ready-to-analyze datasets irrespective of recording conditions and animal species. By simplifying the transformation from raw DeepLabCut outputs to meaningful kinematic insights, refineDLC significantly enhances accessibility and usability, enabling a broader range of researchers—especially those with limited programming expertise—to perform precise quantitative analyses.

Looking forward, adaptive filtering algorithms and real-time quality assessment features are potential enhancements that could further optimize performance, expand pipeline applicability, and automate analysis. Thus, RefineDLC not only addresses the current limitations in markerless tracking technologies but also sets the stage for future advancements in precision phenotyping, behavioral ecology, animal science, and conservation biology.

## Introduction

The DeepLabCut software package revolutionized wide-field behavioral and locomotor studies by enabling precise, markerless pose estimation using deep learning techniques (Mathis et al., 2018). The tool, originally developed to be used in neuroscience and behavioral studies, has also been implemented in the field of animal science and conservation biology (Pereira et al., 2019). By providing an accessible platform for tracking and analyzing movement without the need for the physical contact with the animal, it enhanced the possibilities of conducting behavioral and kinematic studies on big samples as well as on feral populations. To this day, DeepLabCut has been successfully applied in research on numerous species, including horses (Bucci et al., 2025), dogs (Gill et al., 2024), cattle (Zhao et al., 2023) or non-human primates (Wiltshire et al., 2023). However, most of these studies focused on the specific behaviors rather than quantitative kinematic traits and did not develop a robust pipeline for further quantitative analysis.

Therefore, despite its widespread adoption and the availability of an accessible Graphical User Interface (GUI), the usability of DeepLabCut is somewhat limited by the computational expertise required to interpret and refine the outputs. The raw bodypart coordinate data can be noisy and challenging to process, especially when the accuracy of the model is low. Translating these raw outputs into meaningful kinematic parameters often requires additional processing steps, which can be a barrier for researchers without a strong background in programming and data analysis.

Thus, there is a need for a comprehensible and user-friendly post-processing pipeline to enhance the usability of DeepLabCut. Such a pipeline should streamline the transformation of raw coordinate data, limiting noisiness, as well as false positive and false negative labelling. By simplifying data cleaning, filtering and interpolation, a standardized post-processing framework can facilitate more accurate and efficient analyses.

This paper addresses this need by presenting an accessible post-processing pipeline designed to facilitate kinematic analysis from DeepLabCut outputs. We aim to provide researchers with a tool that bridges the gap between raw noisy outputs and ready-to-interpret datasets, enhancing the overall utility of DeepLabCut in locomotor studies and beyond.

## Methods

### Material

We gathered two distinct datasets: a small set of 17 videos of walking dairy cows recorded in a controlled environment (MP4, 1080p), and large set of 524 videos of trotting horses obtained from YouTube (AVI, 720p) (www.youtube.com/arabian-essence, accessed September – October 2023). The videos of horses were not recorded for experimental purposes and therefore contained many issues such as partial occlusion of the animal, inconsistent camera panning, low video quality etc. Both sets of videos were labeled with a custom-trained resnet-50 model with 1M+ iterations, 2041 training frames for horse dataset and 217 for cow dataset, with the same 22 body parts labeled in both models. In the cow model, train error reached 1.52 px with test error = 6.68 px after 1.03M training iterations. In the horse model, after 1M iterations, train error was 8.77 px while test error = 12.68 px.

### Data processing overview

Our data processing pipeline begins with essential data cleaning steps to prepare the output from DeepLabCut for further analysis. These preprocessing procedures address specific issues that previously identified during analysis of DLC output files: inverting the values of y coordinates so the motion function can be analyzed at two-dimensional Cartesian system, deleting non-processed rows, and discarding body part labels that are not relevant for further (specific) analysis. These steps are followed by filtering of the coordinate values based on the position changes of each labeled point and the likelihood assigned by the DeepLabCut model, and subsequent interpolation to replace removed values. The full workflow is presented in Figure 2. Below, we describe in detail each of the workflow steps and their implementation in the validation datasets.

**Figure 1:**
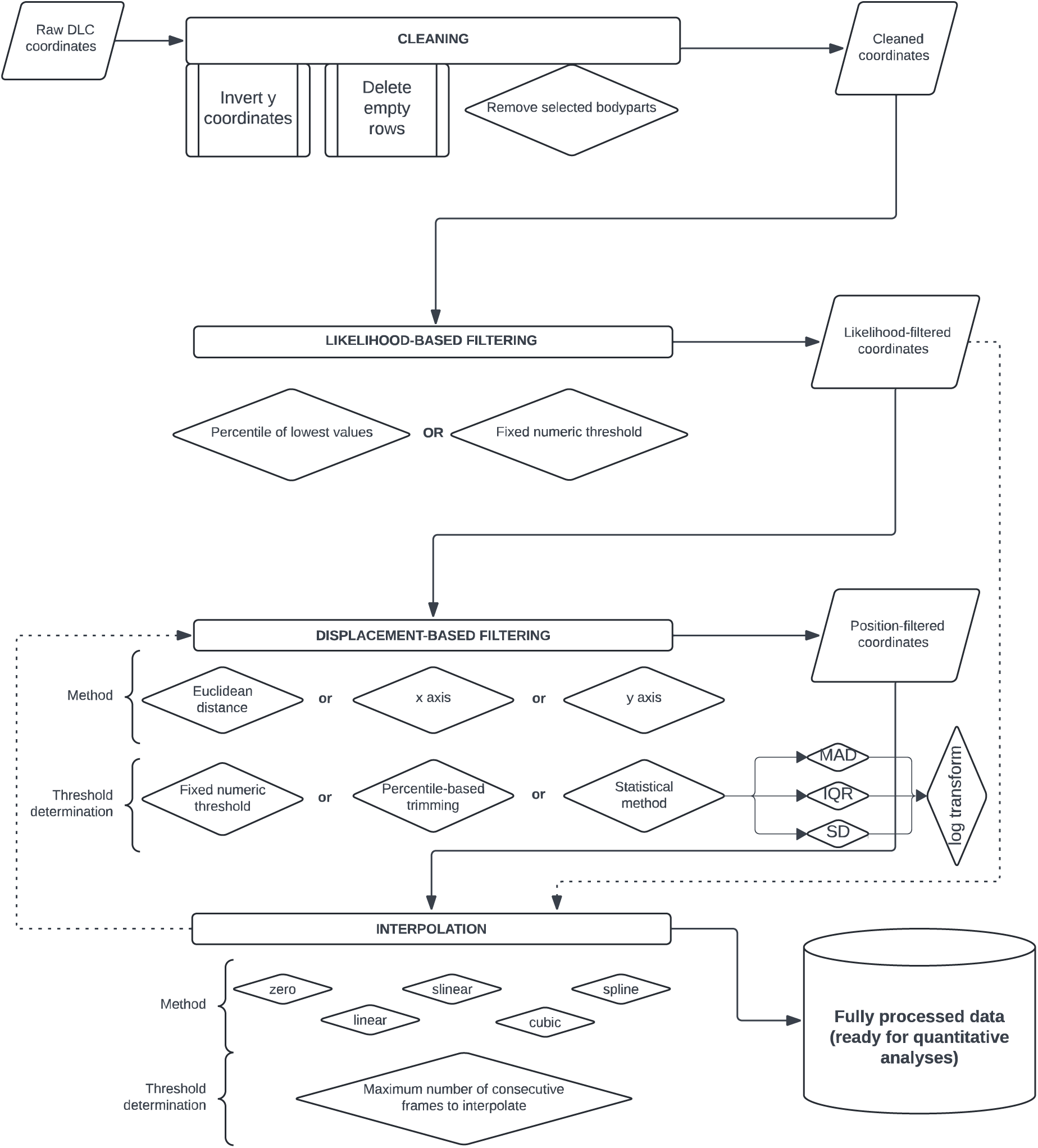
The workflow presented in this study. The assumed input is the raw coordinates obtained with DeepLabCut analysis contained in one directory. The final output retains the input data structure.

**Figure 2:**
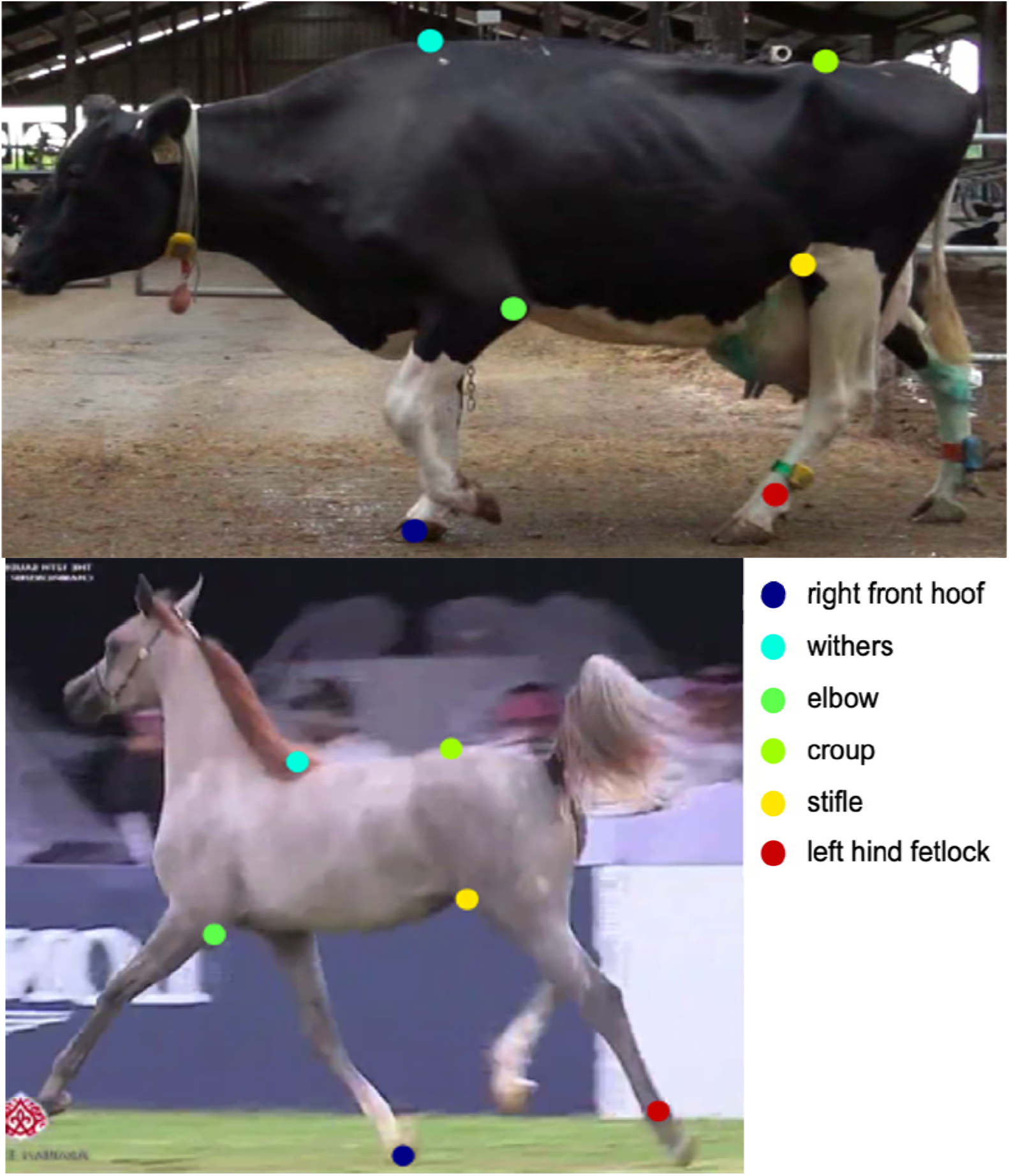
The six body parts selected for pilot analysis and visualization in this study: withers, croup, elbow, stifle, right front hoof, and left hind fetlock.

### Data Cleaning Steps

#### Inversion of Y Coordinates

In computer vision and image processing, the coordinate system of digital images typically places the origin point (0,0) at the top-left corner of the image. In this system, the x-coordinate increases to the right, and the y-coordinate increases downward. This convention stems from the way images are stored in memory and displayed on screens, where pixels are mapped starting from the top-left corner and moving down row by row. DeepLabCut follows this standard image coordinate system when outputting the tracked positions, resulting in y-coordinate values that increase as the position moves downward in the frame. However, for many analyses, especially those involving physical movement in real-world space, it is more intuitive and conventional to have a coordinate system where the origin is at the bottom-left, and the y-coordinate increases upward. For instance, when a limb moves upwards, the y coordinate value increases instead of decreasing.

To align the tracking data with a more traditional Cartesian coordinate system and facilitate intuitive interpretation of vertical movements (where upward movement corresponds to increasing y-values), we included the option of inverting the y-coordinate values in the dataset. This transformation effectively mirrors the y-axis, reorienting the coordinate system so that higher y-values represent positions higher in the frame. By doing so, analyses that rely on vertical displacement—such as calculating the height of jumps or any other upward movements—become more straightforward and align with conventional mathematical representations.

#### Removal of Zero-Value Rows

During processing the horse dataset (videos in AVI format), we observed that some frames contained zero values across all coordinates and body parts (always in the end of the video). This issue can be caused by corrupted video frames, variable frame rates, or video encoding issues. This anomaly also arises from an occasional error within the DeepLabCut processing pipeline, which has yet to be fully addressed by the software developers. These zero-value rows do not represent valid tracking data and can introduce significant inaccuracies, especially when calculating descriptive statistics, if left uncorrected. The issue can be resolved by re-encoding the video using H.264 codec with a constant frame rate and standard pixel format with tools such as FFmpeg or converting video format from AVI to MP4. However, if the rate of the non-analyzed frames is small, the simplest solution can be to remove the zero values for non-processed frames. We implemented a procedure to identify and remove any rows where all coordinate values are zero. By excluding these erroneous data points, we enhance the overall integrity of the dataset and prevent potential distortions in the analysis. Additionally, we implemented a function that outputs the total number of frames, and the number of frames corrupted in every file of the dataset to facilitate calculation of summary statistics for this issue.

#### Exclusion of selected body parts

In certain cases, it is helpful to construct a model that includes body parts that are not relevant to the analysis but are helpful for improving the model performance. For instance, in the horse model, the handler’s legs were annotated to help the model distinguish them from the animal’s legs, reducing misidentification during tracking. Yet, data corresponding to the handler’s legs are irrelevant for analyzing the animal’s movement patterns. In other cases researchers may want to focus on just a subset of previously annotated keypoints in a specific subanalysis. Therefore, we included the option of removing indicated body parts from the dataset after initial processing. This step is optional and can be tailored based on the specific requirements of the analysis. By focusing solely on the relevant body parts, we streamline the dataset and, most importantly, reduce computational complexity.

We applied all three cleaning options to both validation datasets, discarding 16 from initially marked 22 bodyparts, and focusing on six presented in the Figure 2.

### Data filtering

Following the initial steps of data cleaning, we filtered the data based on various criteria, excluding not only low-likelihood labels, but also capturing “false positive” values. Two main filtering steps are implemented: firstly, we filter out the lowest-likelihood frames, based on the values assigned by DeepLabCut. Keeping in mind that some of the erroneous labels, dubbed “false positives”, may evade the likelihood filter, we then focus on position change-based filtering. This method allows identification of sudden and physiologically improbable label movements, even if the likelihood is high.

#### Likelihood-Based Filtering

Each bodypart label in every frame in the DeepLabCut output is assigned a probability of the correct labeling. Based on the value of this likelihood, the user can filter the output to remove frames with the low likelihood. This can be done by either setting an absolute likelihood threshold, or by defining a proportion (percentage) of the lowest likelihood values to be discarded. Since the likelihood distribution can vary radically between bodyparts, if the ‘percentile’ option in the presented workflow is selected, the distributions of the likelihood values are calculated for each bodypart independently. In the validation process, we employed two strategies. In the first case, we removed 5% of the lowest likelihood assignments for every bodypart in both datasets. In the second case, we applied global cut-off likelihood threshold = 0.5.

#### Displacement-based Filtering

The incorrectly labeled points that slip through the likelihood filter can be further detected and removed based on the displacements, i.e., the changes in their position between neighboring frames. Physiologically improbable values can be removed based either on the Euclidean distance for the same bodypart label in consecutive frames or just on the linear change along x or y axis. The Euclidean-distance-based option is in general more robust. The x or y axis option is useful if the main goal is to calculate some speed-related parameters or during the analysis of vertical movements, such as jumping.

Since the distribution of the data with erroneous displacement values will naturally be right-skewed, we employed statistical methods that are designed for identifying outliers in such distributions. Three main filtering modes are available. In the “absolute threshold” mode, the Euclidean distance or the difference in x- or y-values between consecutive frames are calculated for each body part. If the computed distance exceeds a user-specified fixed threshold – a predefined numeric cutoff (in pixels), frames exceeding this positional displacement threshold are flagged and removed from further analysis. The percentile-based trimming removes (winsorizes) positional changes above a user-defined percentile threshold (e.g., 99^th^ percentile), regardless of their absolute magnitude. Alternatively, the threshold can be determined based on the outliers in the distribution of the position changes, detected with the use of one of the statistical methods: median absolute deviation (MAD), interquartile range (IQR), skew-adjusted IQR (Hubert & Vandervieren, 2008), or standard deviation (SD). Additionally, before robust outlier detection, displacements may optionally be transformed by the logarithmic function, stabilizing variance and reducing skewness in the displacement distributions.

Given the highly right-skewed displacement distributions observed in the validation datasets (Supplementary File 1), we adopted two displacement-based filtering approaches, both based on Euclidean distance. First was based on absolute threshold set to 30 pixels for all bodyparts. This value was determined based on the screening of the trajectory plots. Our second approach used skew-adjusted IQR fences with logarithmic transformation and IQR multiplier = 1.5. Both filters have been applied to the dataset after 5th percentile likelihood filtering. In the case of IQR-based filtering, the “NaN” values resulting from likelihood-based filtering were masked from calculation of the quartiles.

### Interpolation of filtered data

The filtering steps naturally result in missing data points where coordinates values were set to NaN (not a number). To address gaps in the dataset and maintain continuity, the user can interpolate missing data using one of five methods: zero (zero-order hold), linear, slinear (first-order spline,) cubic (third-order spline) or spline (polynomial/B-spline of arbitrary order). The choice of the method depends highly on the type of behavior and movement being analyzed. The summary of the differences between these methods, along with the examples of usage, is presented in Table 1.

**Table 1:**
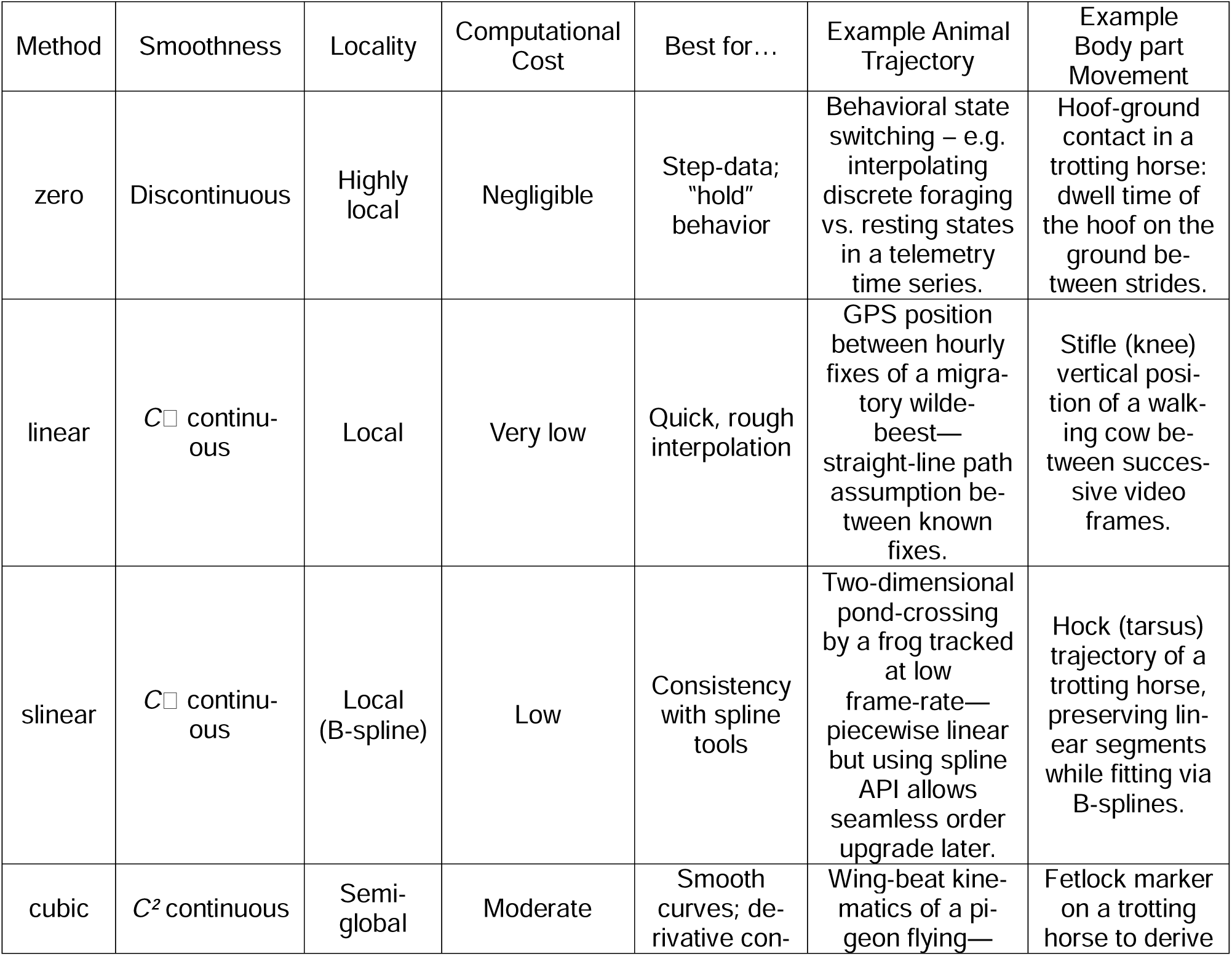
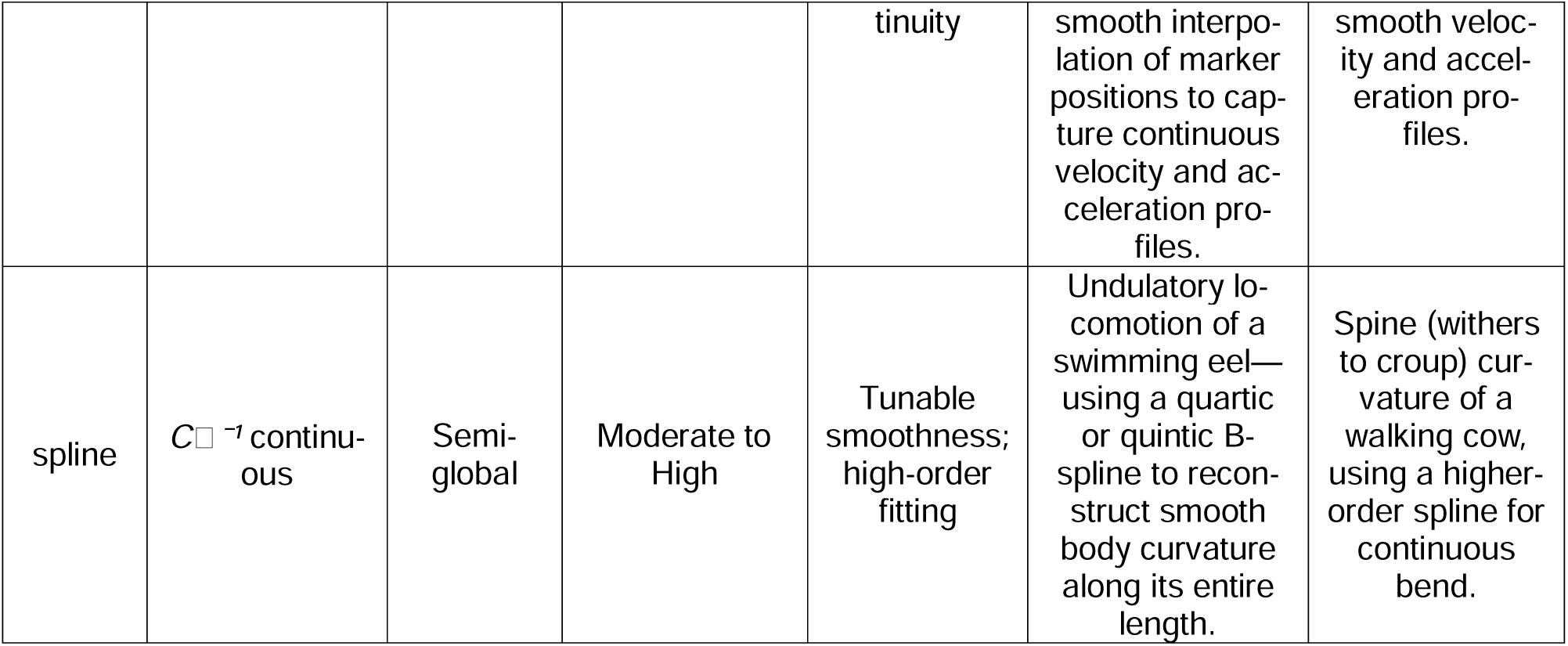
Comparison of interpolation methods available in the presented pipeline, with ilustrative animal-movement trajectories and example body-part movements in walking cows and trotting horses.

The interpolation step concludes with choosing the maximum number of the consecutive missing points that should be interpolated. This depends both on the video recording frequency (which can be translated into seconds, e.g. five frames in 30 FPS video is equal to 0.1 seconds) and the type of the studied locomotion (e.g., the speed and swiftness of the animal). For the validation of the method, we uniformly applied spline interpolation to all bodyparts, with maximum number of the consecutive missing points interpolated of five. The interpolation has been performed on the datasets with filtered 5% lowest values of likelihood and the displacement values higher than 30.

## Software implementation overview

The refineDLC post-processing pipeline is implemented as a modular Python package (Python version 3.10 or higher) consisting of four standalone scripts: clean_coordinates.py, likelihood_filter.py, position_filter.py, and interpolate.py. Each script is designed to be executed via a command-line interface (CLI), facilitating reproducibility, automation, and integration into larger data-processing pipelines.

The first step (clean_coordinates.py) addresses initial data preparation, including inverting y-coordinate values, removing zero-value frames, and optionally discarding irrelevant body parts. Users specify input and output file paths as command-line arguments, along with an optional parameter listing body parts to exclude from further analysis.

Subsequently, the second step (likelihood_filter.py) filters the cleaned data based on user-defined likelihood thresholds or percentile-based criteria. Low-confidence data points identified by DeepLabCut likelihood scores below a fixed threshold or within a user-specified lowest percentile per landmark are replaced with NaN (not a number), effectively excluding unreliable coordinate values. User specifies the input and output directories or file paths along with either the fixed likelihood threshold value or percentile.

The third step (position_filter.py) applies filtering based on positional changes between consecutive frames. Users select from three filtering methods (euclidean, x, or y) and set thresholds using either fixed positional limits, percentile of the highest values, or robust statistical methods (standard deviation, median absolute deviation, or interquartile range). Data points exceeding these criteria are marked as NaN to denote exclusion.

Finally, the interpolate.py script addresses gaps resulting from the previous filtering processes. Users can choose from five interpolation methods (zero, linear, slinear, cubic or spline) and set the maximum allowable gap size for interpolation. This maintains data continuity while still respecting the temporal dynamics of motion. Each script provides detailed logging, facilitating debugging and data tracking. Comprehensive documentation and example command-line usage are included in the repository’s README.md file, promoting ease of adoption and reproducibility across research contexts. The entire refineDLC pipeline, along with installation instructions, usage examples, and source code, is publicly available via GitHub (https://github.com/wer-kle/refineDLC).

## Results

### Data cleaning

The total disk size of the two datasets containing raw CSV outputs from both DeepLabCut models was 75.1 MB (70.7 MB of horse dataset and 4.4 MB of cattle dataset). The cleaning steps reduced this size to 15.1 MB for horse dataset and 1.6 MB for cattle dataset (16.7 MB in total). In 367 out of 524 files in the horse dataset (videos collected in AVI format) we identified at least one “zero row”, with the maximum of 128. In total, 16,059 out of 61,672 rows in the horse dataset were empty. In the cattle dataset, we didn’t identify any “zero rows”.

### Likelihood filtering

The threshold values marking 5% of the lowest-likelihood values varied notably across datasets, bodyparts, and videos. The distributions of the values representing 95% of the likelihood for each bodypart are presented in the Figure 3. The distal limb markers – left hind fetlock and right front hoof – were the least consistent among all bodyparts. In the horse dataset we observed multiple outliers, and in many cases the cut-off value for 5% lowest likelihood values was close to 0.

**Figure 3:**
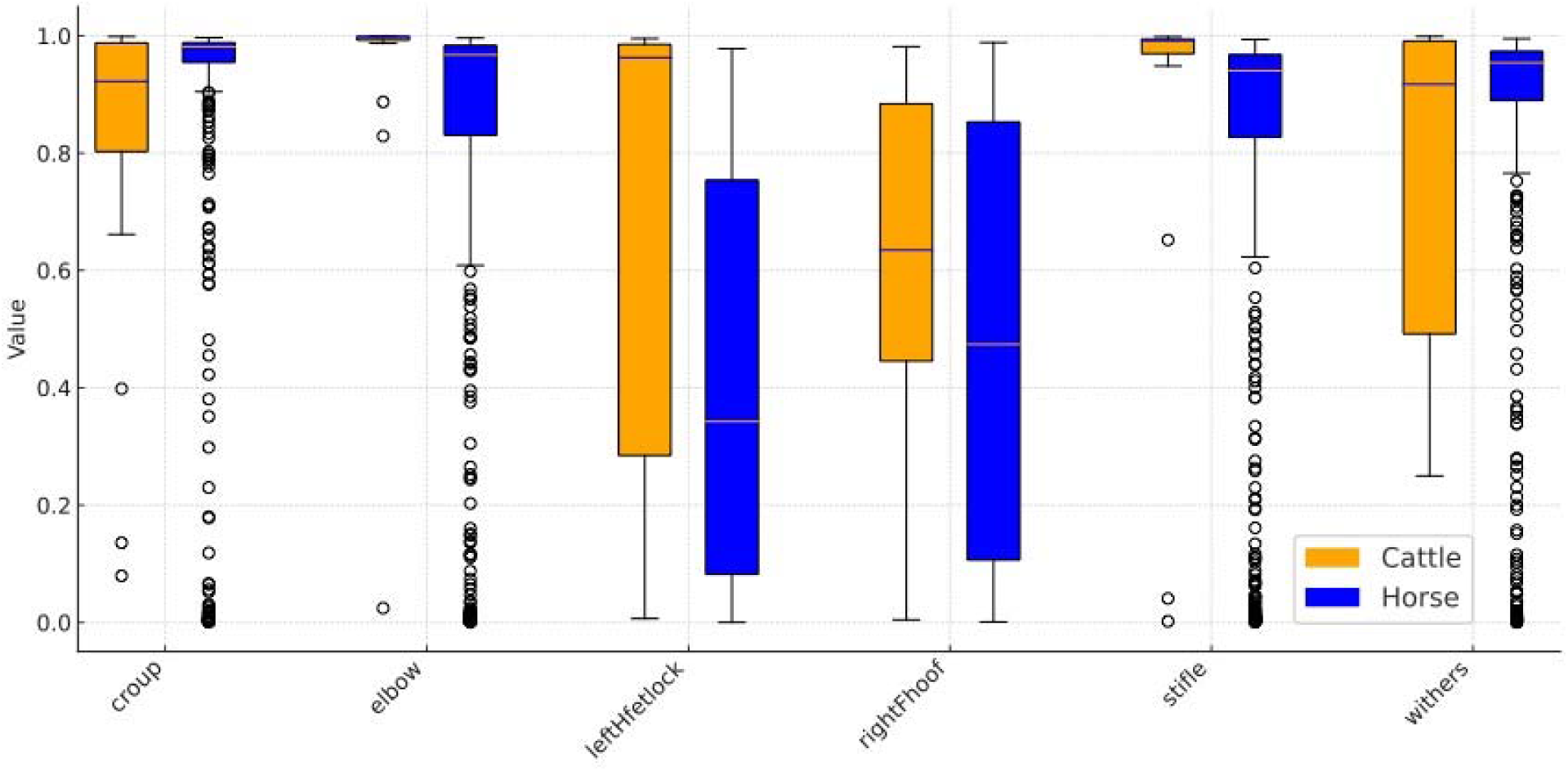
The distribution of cut-off values when with likelihood filter removes 5% of the lowest values. Applying conservative cut-off threshold of 0.5 resulted in non-proportional removal of a notable percentage of frames, surpassing 10% in some cases (Table 2). There were few differences observed between datasets, but, similarly to the first method, distal limb markers (left hind fetlock and right front hoof), had distinctly higher percentage of observations removed (likely due to more frequent errors for these highly mobile keypoints). Moreover, the maximum percentage of frames removed was 89.28 (elbow) for horses and 70.51 (left hind fetlock) for cattle, making the data for these keypoints virtually unusable.

**Table 2:** The average percentage of values removed after applying likelihood threshold = 0.5.

| Bodypart | Cattle Average | Horse Average |
| --- | --- | --- |
| croup | 2.86 | 1.96 |
| elbow | 2.76 | 3.14 |
| left hind fetlock | 10.83 | 10.92 |
| right front hoof | 8.41 | 10.10 |
| stifle | 4.43 | 3.03 |
| withers | 2.92 | 2.28 |

### Displacement-based filtering

The results of displacement-based filter application are presented in Table 3. While the maximum values in IQR-filtered cattle dataset were similar to the arbitrarily selected threshold of 30 pixels, the physiologically impossible values were still left within the horses dataset using this approach (Figure 4).

**Figure 4:**
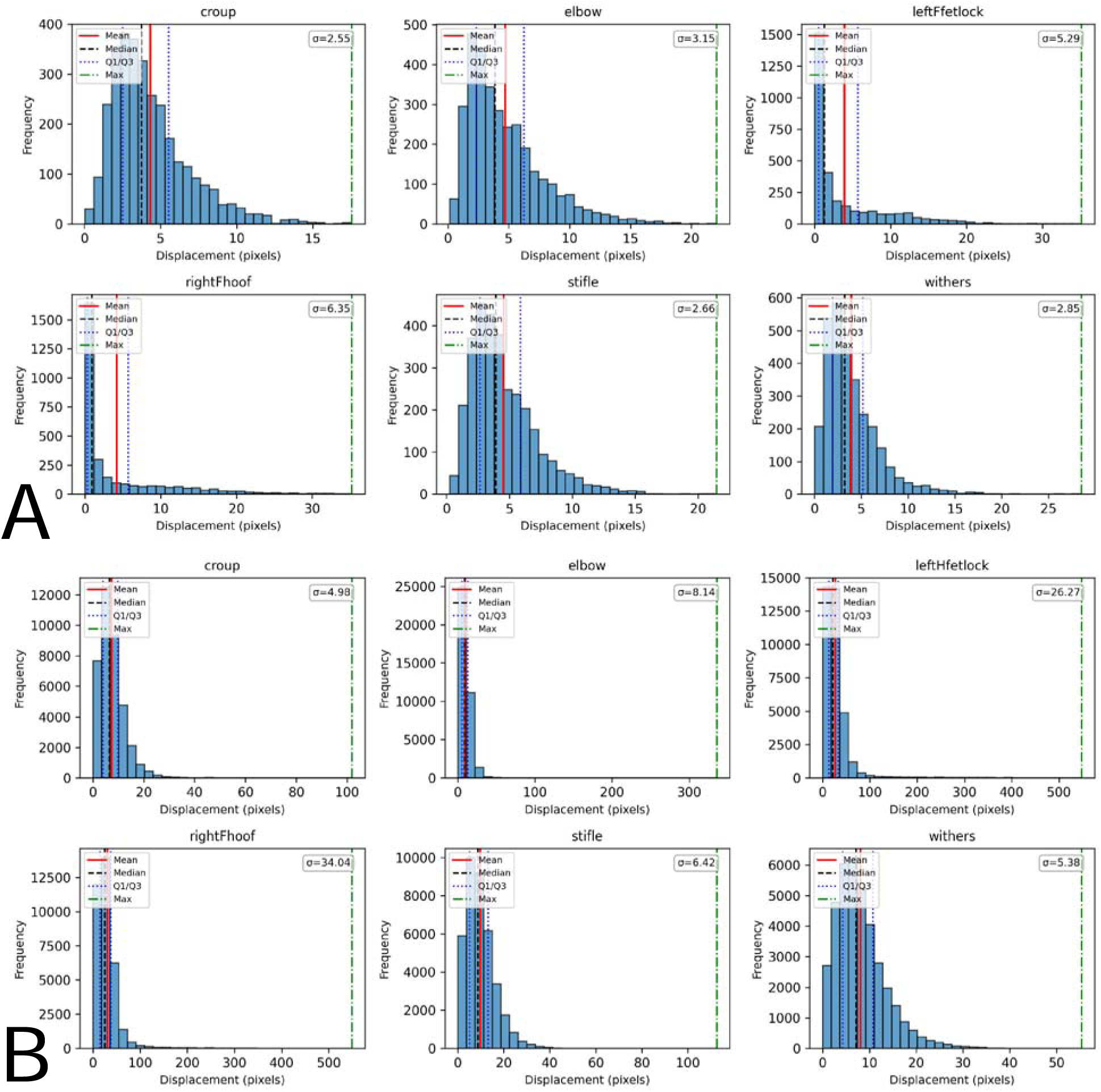
Distributions of the displacement values in cattle (A) and horses (B) datasets after filtering of 5 percentiles of the lowest likelihood assignments and values above 1.5 Interquartile Range (IQR).

**Table 3:** Mean and range of frames removed by two filtering methods—Euclidean distance (>30 px) and IQR-based—across cattle and horse datasets.

| Filtering method | Dataset | Mean number of removed frames | Range of removed frames number |
| --- | --- | --- | --- |
| Euclidean distance<br>(> 30 px) | Cattle | 28.5 | 0 – 190 |
|  | Horses | 134.8 | 0 – 830 |
| IQR-based filter<br>(>1.5 IQR) | Cattle | 95.5 | 16 – 294 |
|  | Horses | 41.3 | 0 – 306 |

### Interpolation and the readability of the locomotion patterns before and after post-processing steps

Application of the interpolation proceedure resulted in replacing 42.8 missing values on average from cattle dataset and 78.9 missing values on average from horse dataset. To compare the readability of the raw coordinates versus the post-processed data, we chose intermediately noisy files (with the number of frames removed by the displacement filter close to the average for the whole dataset), and we plotted the movement trajectories of the bodyparts from both raw (cleaned) coordinates and from interpolated ones (cattle in Figure 5, horses in Supplementary File 2). The application of the filtering and interpolation techniques described above resulted in the uncovering of the easily understandable and interpretable movement trajectories that can be further analyzed in a kinematic or behavioral study.

**Figure 5:**
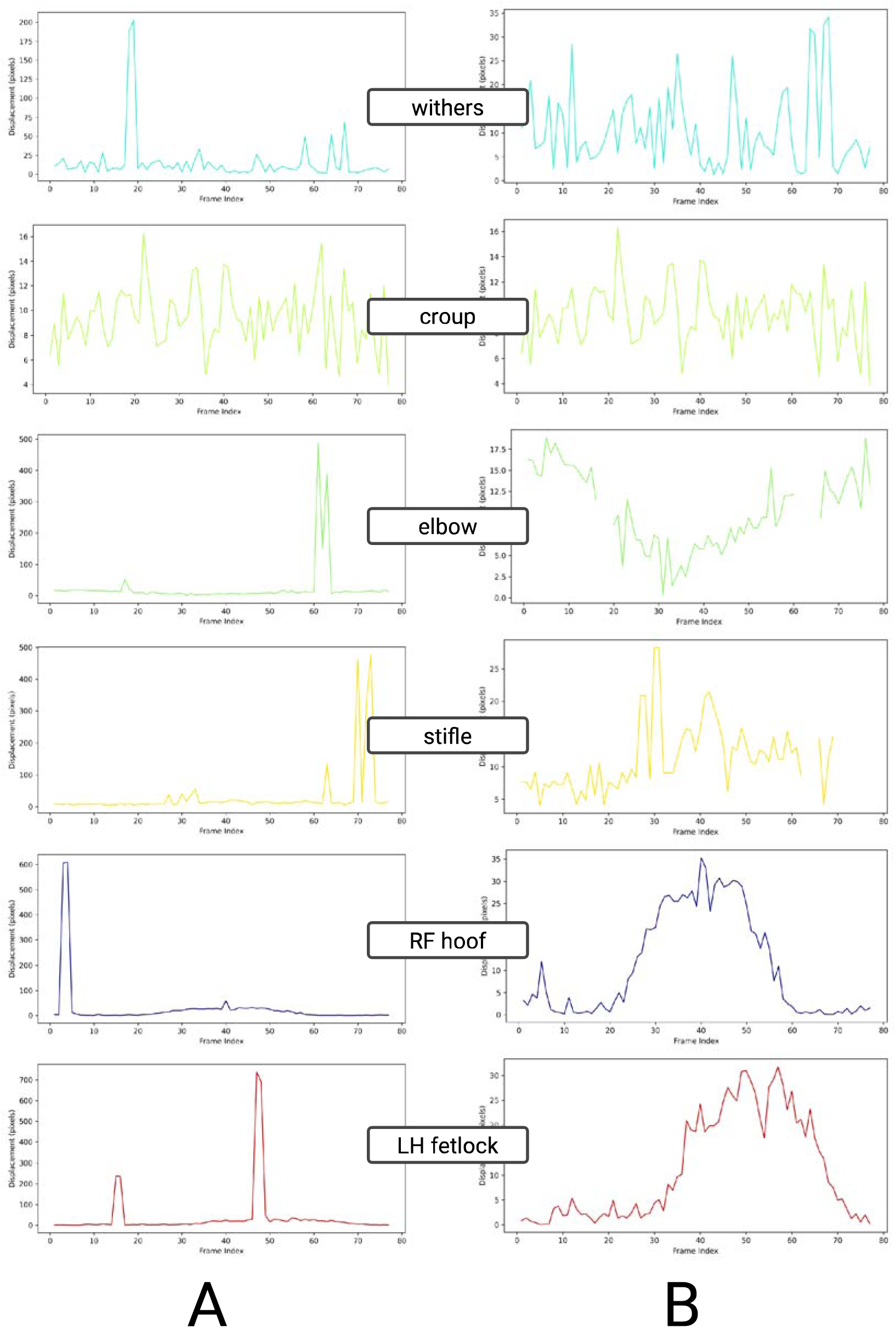
Movement trajectories, presented as bodyparts displacements, before (A) and after (B) the application of the described post-processing pipeline in a selected file from cattle dataset.

## Discussion

The refineDLC post-processing pipeline presented in this study addresses a critical gap in markerless motion tracking by transforming raw DeepLabCut output into robust, reliable kinematic data ready for analysis. Previous applications of DeepLabCut, while groundbreaking in enabling markerless pose estimation (Mathis et al., 2018; Nath et al., 2019), often required substantial computational expertise and labor-intensive post-processing to manage noise from occlusions and low-confidence predictions (Pereira et al., 2019). By contrast, refineDLC provides an intuitive and versatile framework that streamlines this refinement process, effectively bridging the gap and greatly enhancing the accessibility of precise quantitative analyses in behavioral and locomotion studies.

A key strength of refineDLC lies in its sequential data cleaning, filtering, and interpolation steps that collectively refine DeepLabCut outputs. The initial data cleaning stage (including inversion of y-coordinates) corrects a fundamental inconsistency between standard image coordinate conventions and the Cartesian system used in biomechanical analyses, immediately improving the interpretability of vertical movement data. Moreover, the systematic removal of spurious zero-value frames and irrelevant bodypart labels at the outset ensures that only valid, meaningful data points enter subsequent analysis, thereby enhancing the integrity and efficiency of all downstream processing.

In the filtering stage, our pipeline advances beyond traditional likelihood-only filtering approaches. Prior DeepLabCut workflows predominantly relied on dropping low-confidence predictions based on model-assigned likelihood scores (Hayakawa et al., 2024; Wiltshire et al., 2023). However, exclusive dependence on likelihood can be problematic: as we observed, likelihood value distributions vary widely between videos, making any single confidence cutoff difficult to generalize, and a strict likelihood threshold can unintentionally discard large portions of data. More critically, a likelihood-only filter may fail to remove implausible outliers that the model misclassifies with high confidence. For example, sudden non-physiological jumps in a tracked keypoint’s position can still receive high likelihood scores, yielding high-likelihood false positives that evade removal. Our pipeline mitigates these issues by combining likelihood-based filtering with an additional displacement-based filter that examines frame-to-frame positional changes. This dual strategy catches both low-confidence points and anomalously large movements, substantially increasing the robustness of the cleaned data.

Notably, we found the optimal displacement filtering strategy to be dataset-specific: a fixed cutoff of 30 pixels yielded superior results for the cattle dataset, whereas a robust IQR-based filter (using 1.5× the interquartile range) was more effective for the horse dataset. Using a fixed 30 px threshold provides a straightforward criterion to eliminate extreme outliers and proved highly effective in the cattle videos recorded under controlled conditions, where any inter-frame movement exceeding ~30 px was clearly artifactual. In contrast, the IQR-based filter (Hubert & Vandervieren, 2008), which adapts to the variability of each video, was better suited for the trotting horse videos that exhibited erratic camera motion and greater natural variability. By scaling the cutoff to each sequence’s displacement distribution, the IQR method filtered outliers in the horse dataset more contextually, avoiding the over-removal of data that a one-size-fits-all threshold might cause. Indeed, in the heterogeneous horse data, a global 30 px rule tended to remove a large number of frames – including some likely valid points during rapid locomotion or camera shifts – whereas the adaptive IQR criterion preserved more data by only flagging points well outside the normal motion range. However, this adaptiveness has trade-offs: the IQR filter, while conservative, can fail to catch a few truly aberrant points if the overall displacement distribution is heavily skewed. Conversely, the fixed threshold will reliably excise any extreme displacement but cannot distinguish a genuine sudden movement from a tracking error.

Importantly, whatever filtering approach is used, refineDLC’s interpolation step allows the recovery of continuous trajectories from filtered data. Fixed numerical thresholds carry the risk of removing legitimate data when an animal’s movement or a camera jerk produces a large but real displacement. In our pipeline, those removed frames are filled via interpolation – meaning that even if a valid point is temporarily excluded for exceeding a cutoff, its trajectory can be smoothly reconstructed from neighboring frames. This interpolation safeguard helps maintain data continuity and can reintroduce plausible values for points initially deemed outliers. The trade-off is that interpolation may attenuate genuine extreme motions: for instance, a swift limb movement flagged as an outlier will be brought back as a smoothed curve, potentially underestimating its true peak. In this study, we uniformly applied a spline-based interpolation to all body parts (with a five-frame gap limit) based on visual assessments of trajectory quality, ensuring that the reconstructed kinematic profiles preserved the natural temporal dynamics of movement (Skjelbred & Kong, 2019). The combination of judicious filtering and interpolation yielded cleaned datasets in which underlying locomotion patterns were far more discernible than in the raw data.

The usability and effectiveness of refineDLC were demonstrated on two markedly different validation datasets: a controlled-environment cattle locomotion set and a noisy, unconstrained horse trotting set. Despite the stark differences in video quality and recording conditions, the pipeline improved data reliability and clarity in both cases. In the cattle videos – filmed under stable conditions with a static camera and minimal background distractions – refineDLC required relatively mild intervention. The majority of frames had high model confidence, and only occasional outliers (e.g. brief tracking glitches or implausible jumps) needed to be removed. The fixed 30 px displacement filter was sufficient to flag these anomalies, and the resulting trajectories closely reflected the true movements of the cows with minimal interpolation needed.

By contrast, the horse videos from field recordings were characterized by numerous erratic coordinate jumps due to moving cameras, occlusions, and variable lighting. Here, refineDLC’s more aggressive cleaning (particularly the displacement filtering) was critical to isolate genuine motion signals from pervasive noise. The adaptive IQR-based filter proved especially useful for the horses, automatically calibrating to each video’s motion scale and flagging outlying points accordingly. Through this adjusted filtering strategy, refineDLC was able to salvage meaningful trajectories from data that were initially highly inconsistent. These results underscore the pipeline’s versatility and ease of use: with minimal parameter tuning, a user can obtain reliable kinematic data from both well-controlled videos and chaotic, real-world footage. Moreover, the improvements observed across both datasets align with the broader trend in markerless tracking research emphasizing generalizability across diverse conditions (Pereira et al., 2022). In fact, refineDLC’s strong performance on the challenging horse dataset suggests that our approach can offer analysis quality comparable to or exceeding that of alternative pose-tracking systems even in unconstrained environments.

While refineDLC markedly improves the post-processing of DeepLabCut outputs, certain considerations warrant further investigation. Our current filtering thresholds (the likelihood cutoff and the 30 px displacement limit) were chosen empirically based on initial data screening; an important next step is to develop adaptive thresholding strategies that can adjust these values dynamically for different species, movement types, or video conditions. For example, future work could employ machine-learning techniques to automatically optimize the likelihood and displacement criteria, reducing the need for manual tuning. Such an adaptive system might decide when a fixed threshold should be tightened or relaxed or when to prefer a distribution-based criterion, thereby further minimizing user intervention. Additionally, integrating more advanced trajectory-modeling algorithms (e.g., recurrent neural networks or reinforcement learning models) may offer further refinements in distinguishing true movements from artifacts and in guiding the interpolation of missing data.

In conclusion, refineDLC significantly enhances both the usability and the analytic value of DeepLabCut outputs by providing a standardized, user-friendly post-processing pipeline for precise kinematic analysis. By seamlessly integrating data cleaning, dual-mode filtering, and flexible interpolation, the pipeline transforms raw pose-tracking data into high-quality motion trajectories with minimal user effort. This advancement empowers researchers – including those with limited programming experience – to extract meaningful biomechanical insights from complex video datasets. Looking ahead, incorporating adaptive algorithms and additional automation features will further improve refineDLC’s performance and broaden its applicability, helping to bring reliable markerless motion analysis to an even wider range of species, behaviors, and experimental settings.

