## Supplementary File 1 for "refineDLC: an advanced post-processing pipeline for DeepLabCut outputs"

### Distributions of the displacement values

#### Cattle

##### Cleaned

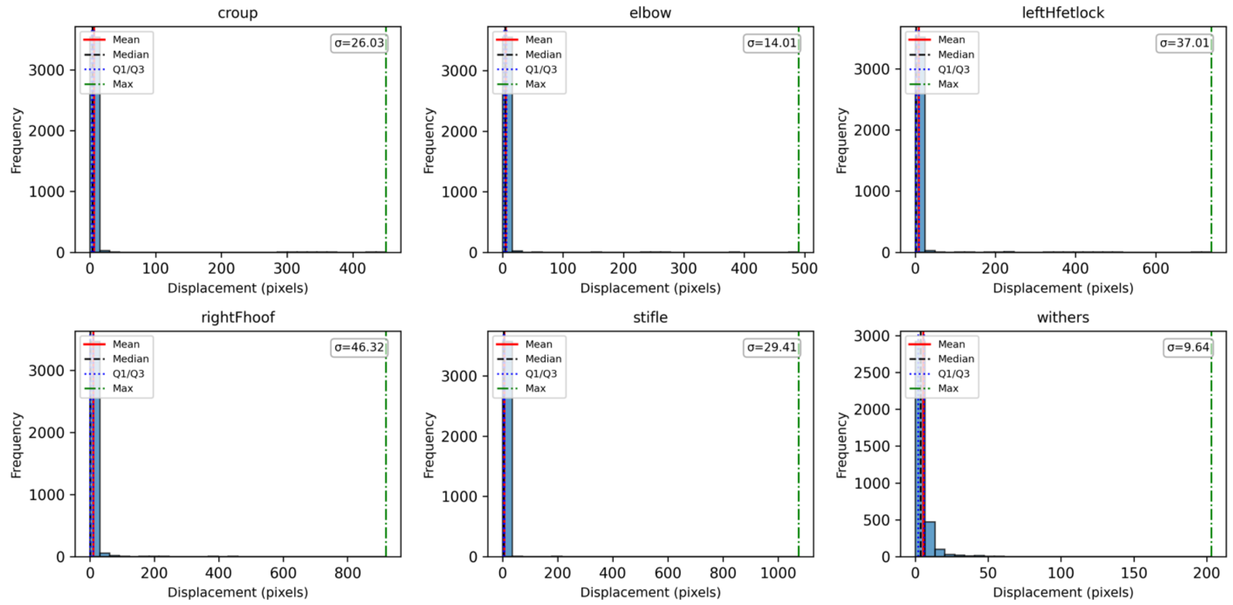


##### Likelihood 5%

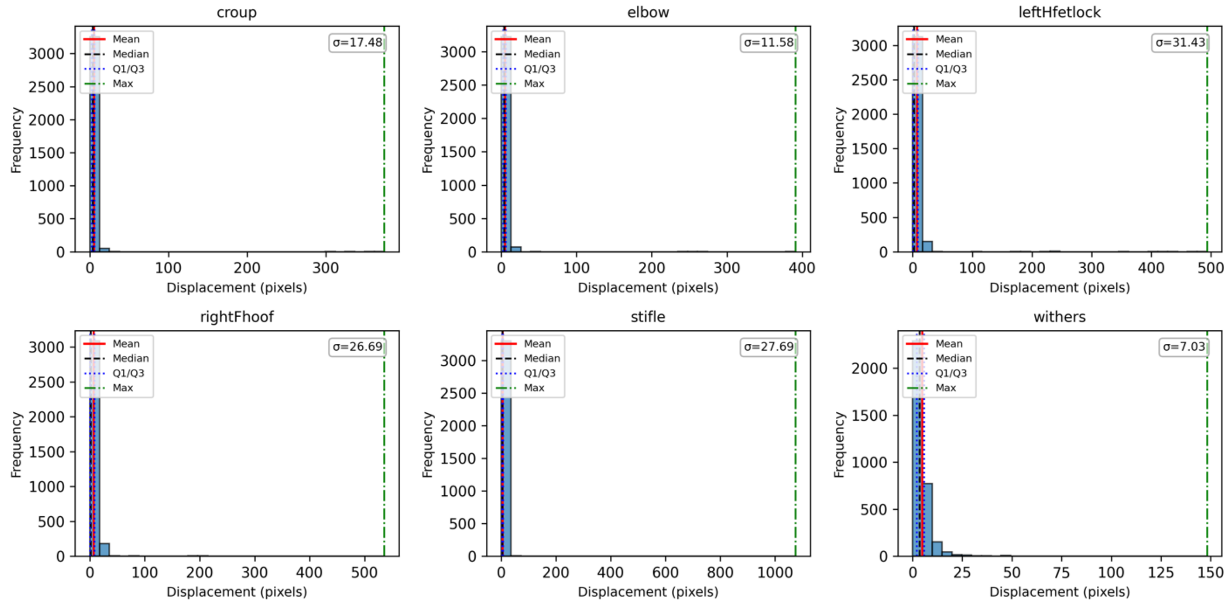

Likelihood threshold 0.5

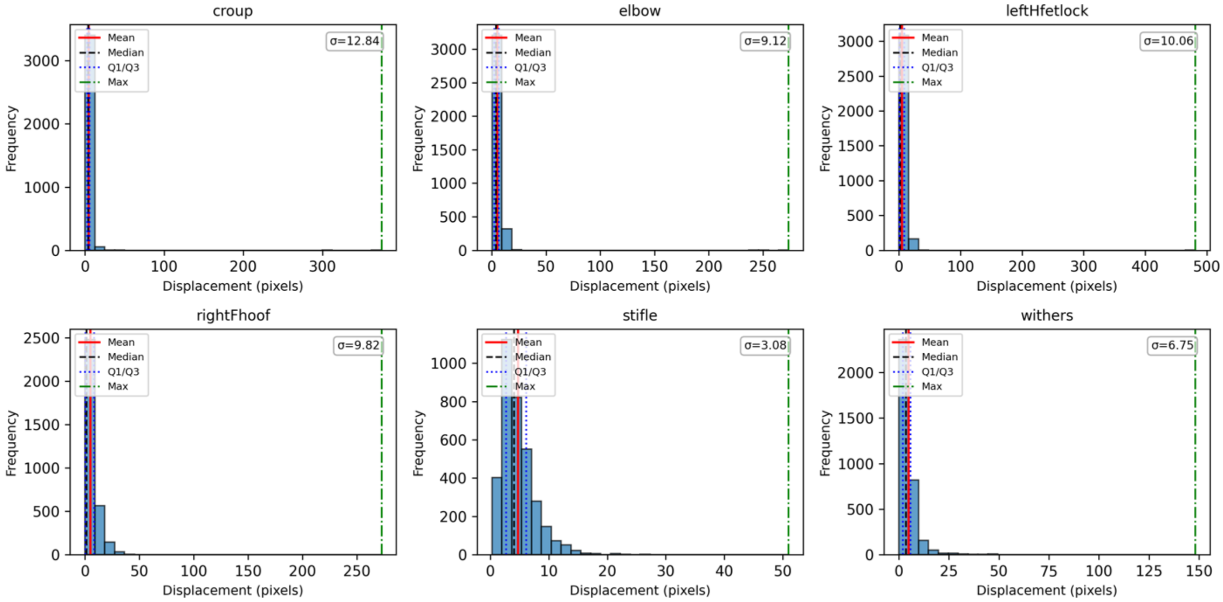


Displacement threshold 30

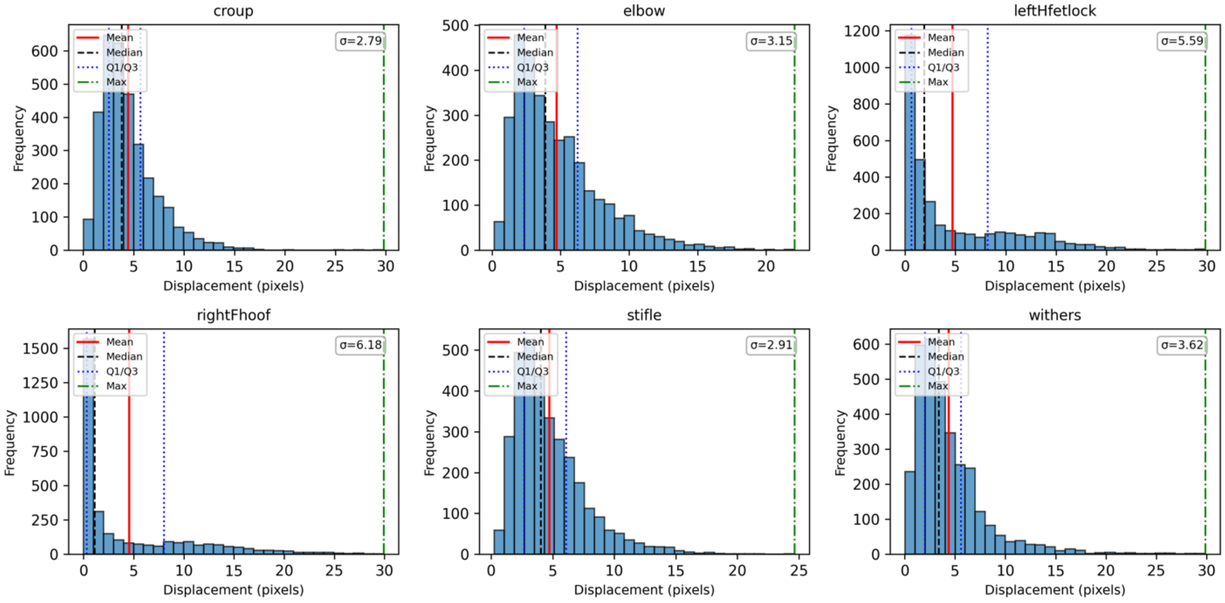

### Displacement iqr 1.5

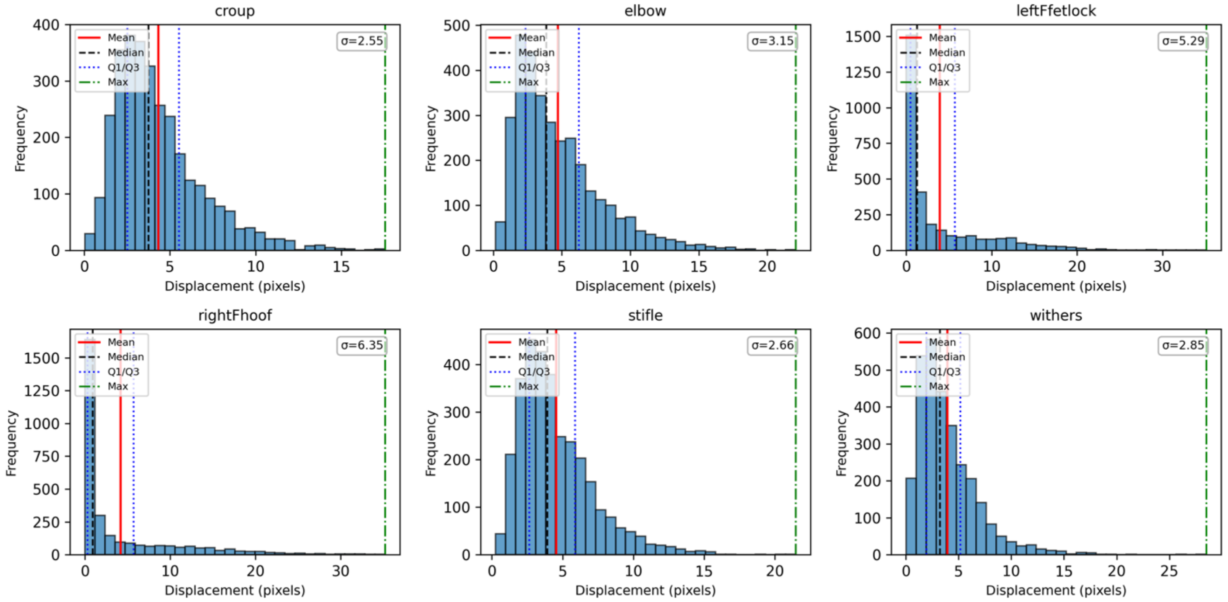


### Interpolated

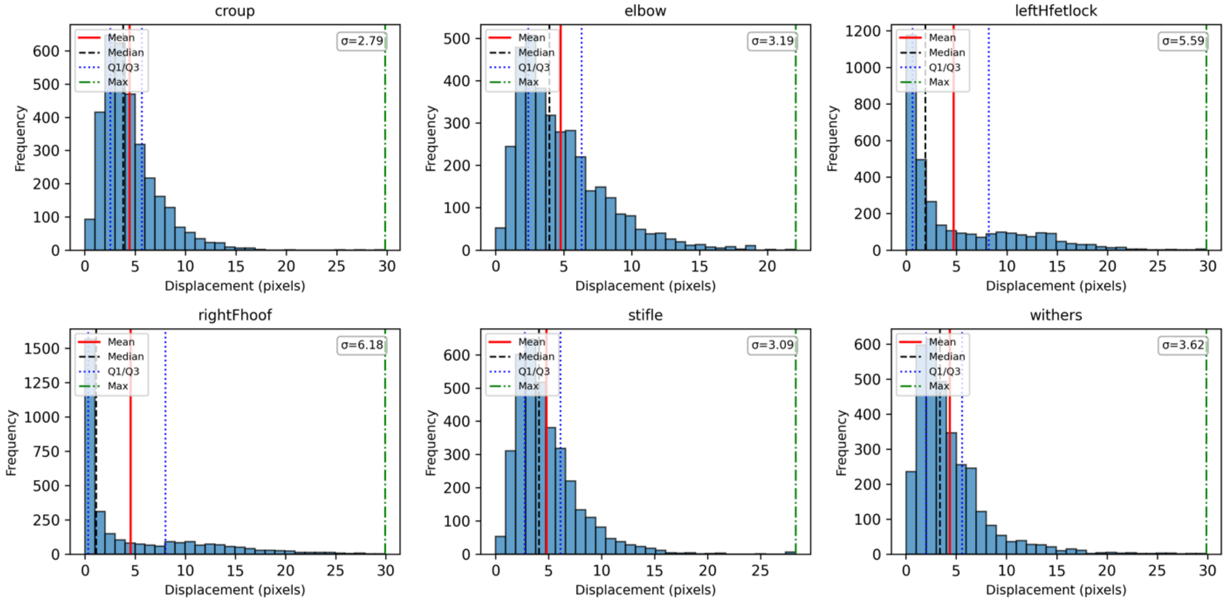

### Horses

#### Cleaned

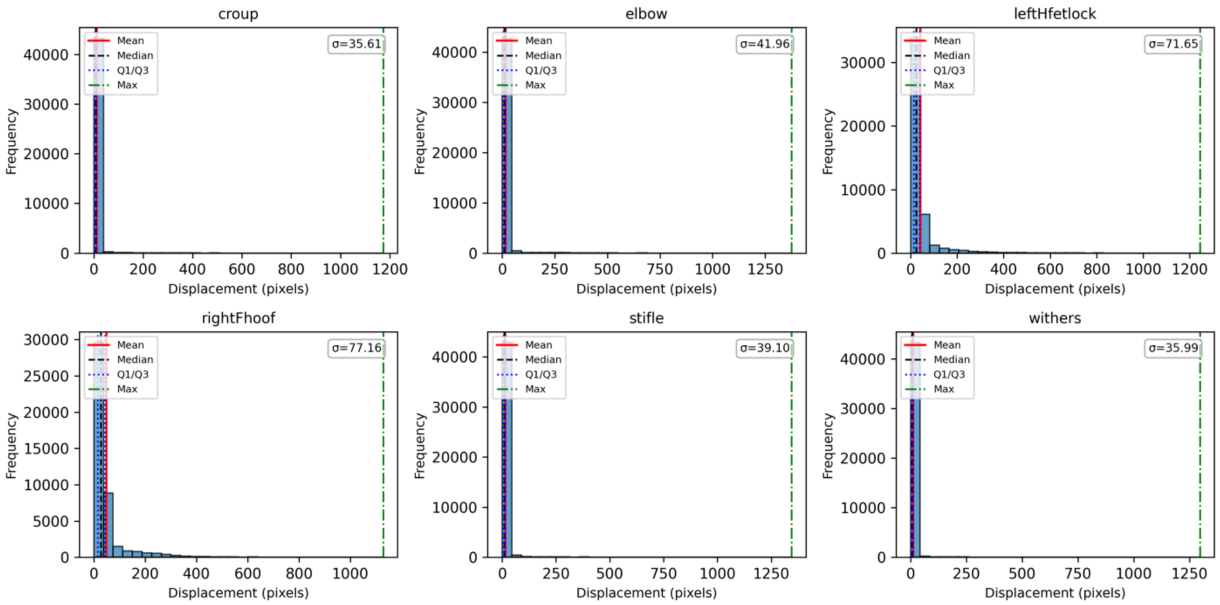


#### Likelihood 5%

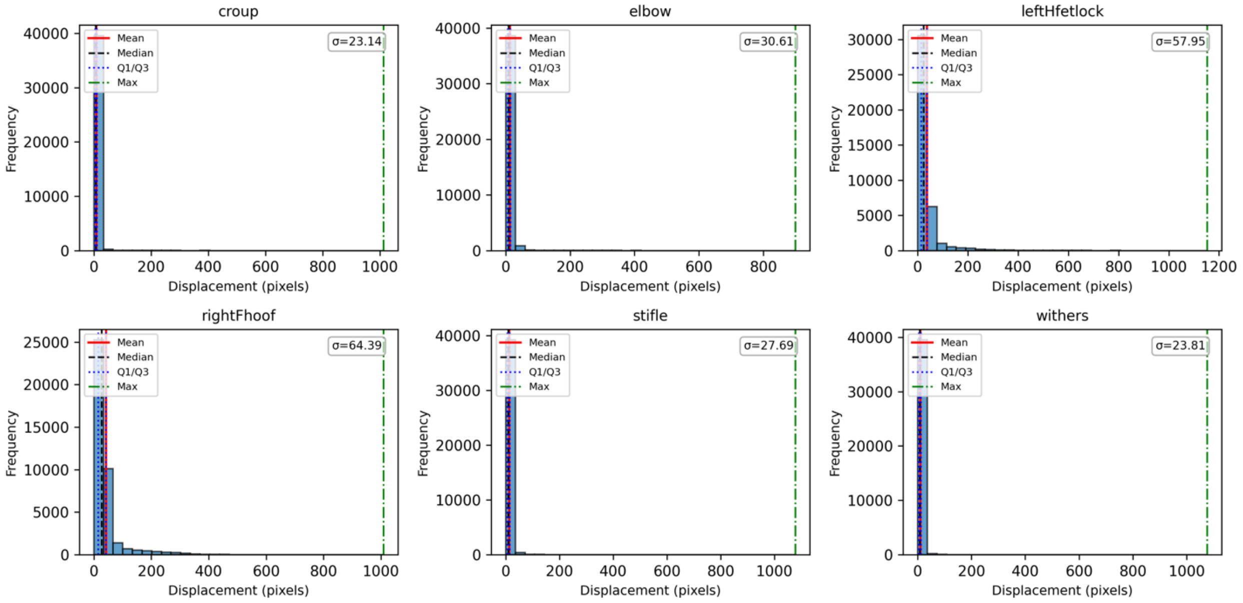

#### Likelihood threshold 0.5

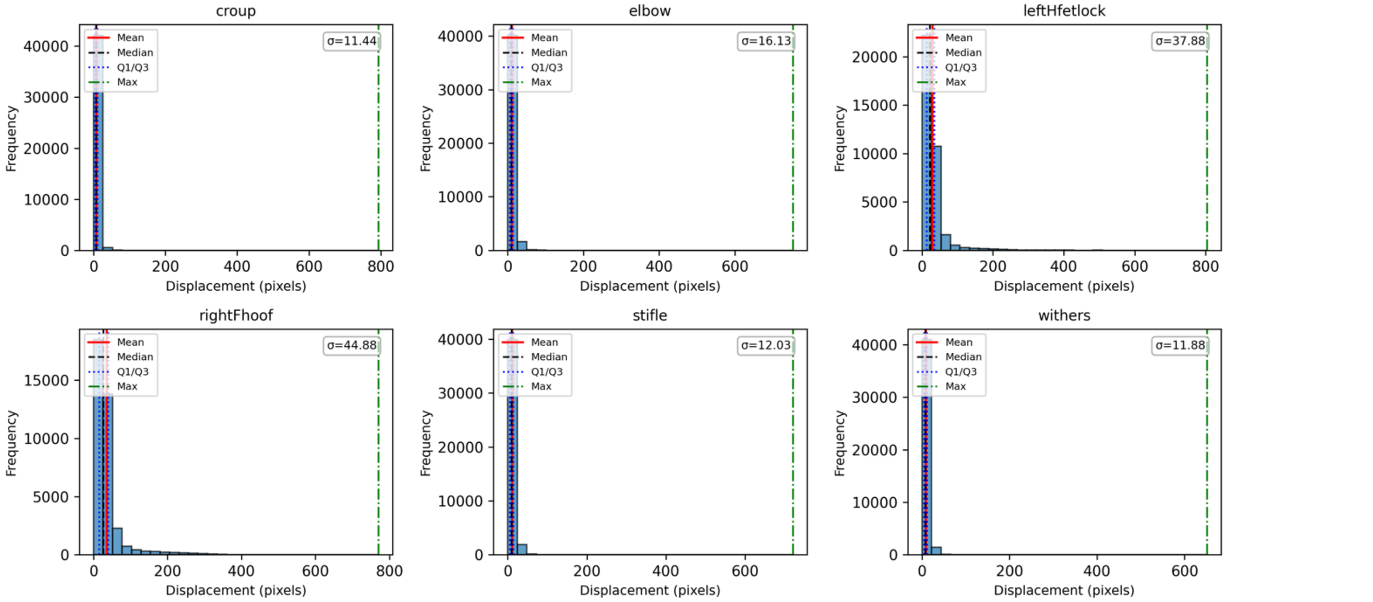


#### Displacement threshold 30

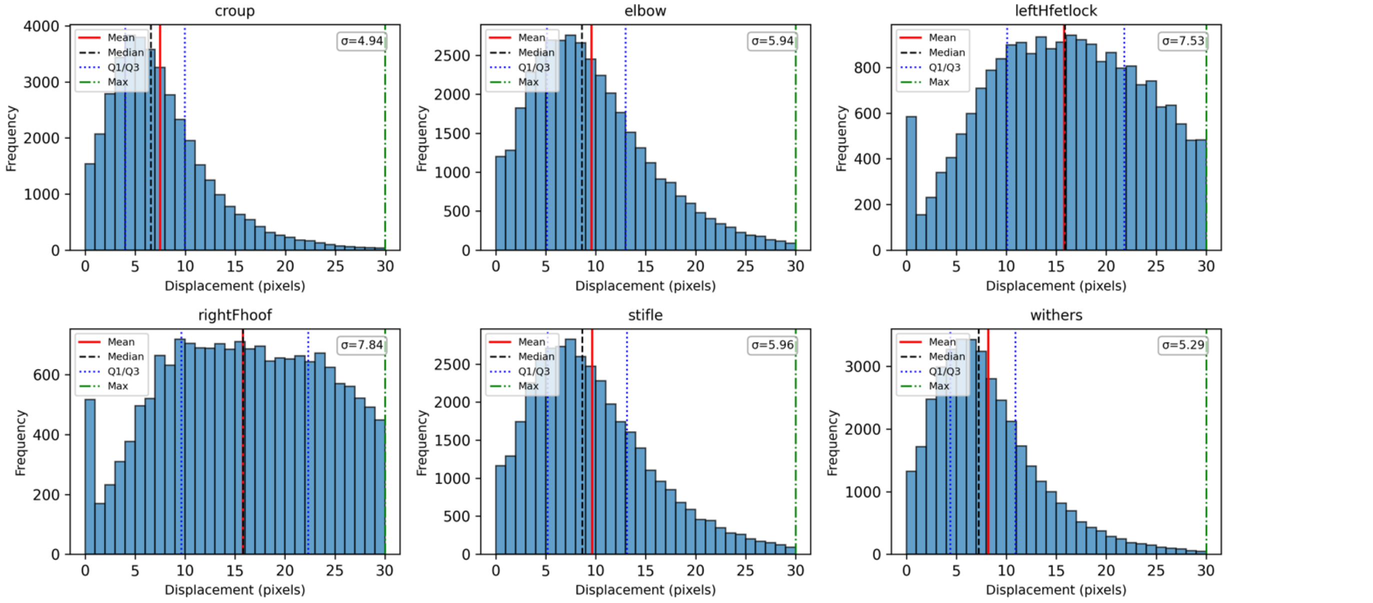

### Displacement iqr 1.5

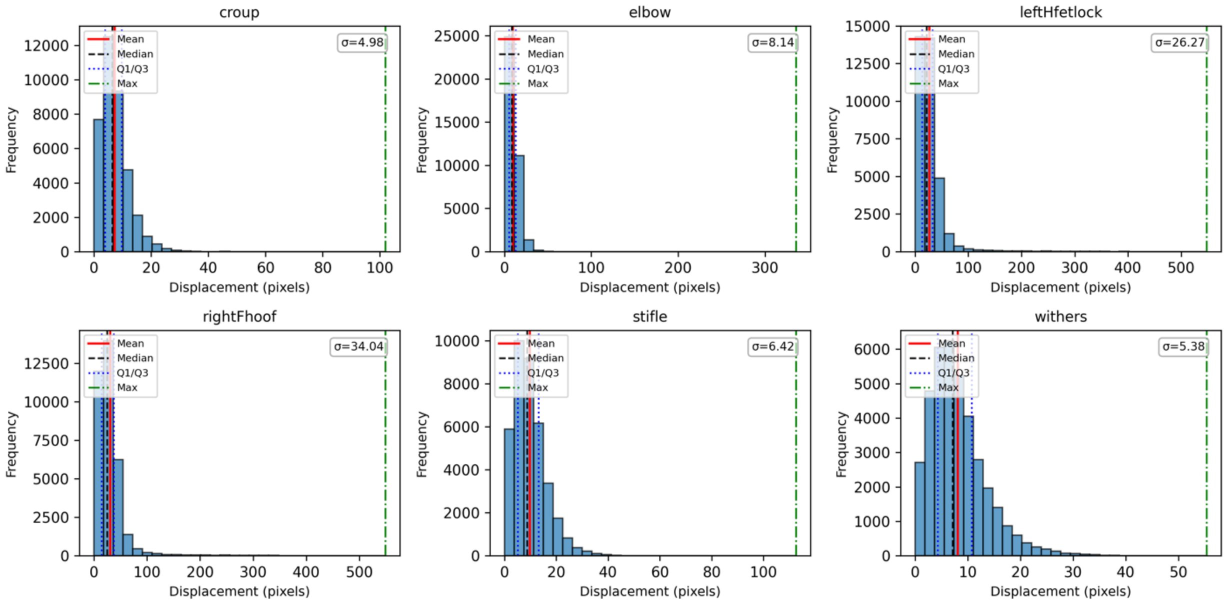
