## Supplementary figures and images for "refineDLC: an advanced post-processing pipeline for DeepLabCut outputs"

### Supplementary File 2

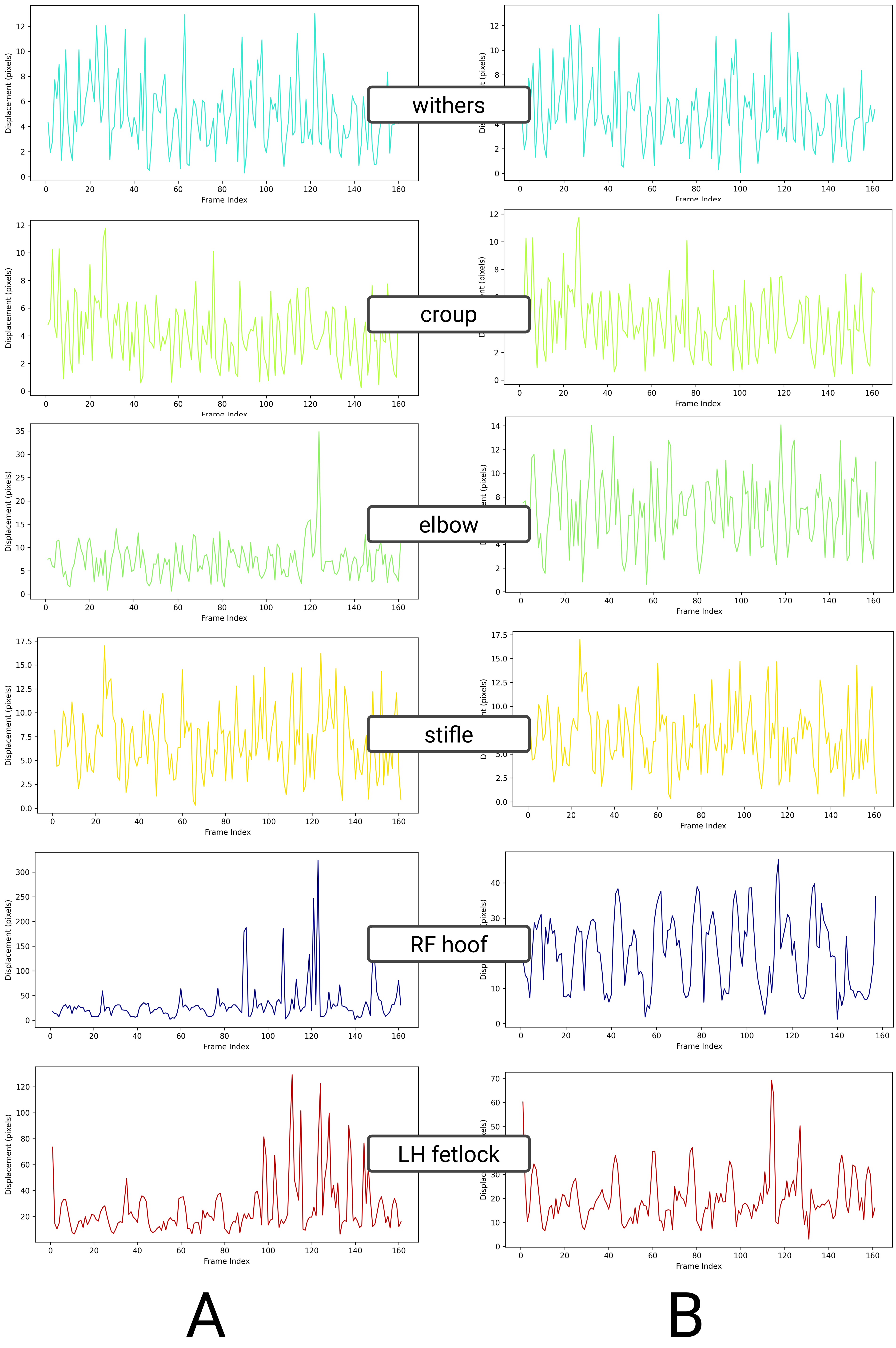
